# Quantitative super-resolution imaging with deep learning segmentation reveals early mitochondrial network remodeling and altered mitochondria-lysosome interactions in Pompe disease muscle

**DOI:** 10.64898/2026.07.21.739567

**Authors:** Ibrahim Hassani, Johan Deniaud, Chantal Thorin, Tony Fiore, Laurence Dubreil, Karl Rouger, Marie-Anne Colle

**Affiliations:** Oniris, INRAE, PAnTher, 44300 Nantes, France; Nikon Healthcare France, succursale Nikon Europe B.V., 94500, Champigny-sur-Marne, France

**Keywords:** Pompe disease, mitochondria, lysosome, super-resolution, image analysis

## Abstract

Pompe disease (glycogen storage disease type II) is an autosomal recessive lysosomal storage disorder caused by acid α-glucosidase deficiency, leading to lysosomal glycogen accumulation, autophagosome buildup, and defective autophagic flux. Mitochondrial abnormalities, traditionally described by electron microscopy as paracrystalline inclusions, cristae loss, and swollen structures, have long been considered secondary consequences of lysosomal dysfunction. However, the spatial organization, organelle crosstalk, and fiber-type-specific progression of these alterations in skeletal muscle remain poorly understood. We combined super-resolution image scanning microscopy with deep learning-based segmentation to quantitatively assess mitochondrial and lysosomal remodeling, and their direct physical interactions, in Pompe disease (*Gaa*⁻/⁻) mice. Organelles were analyzed at pre-symptomatic (1 month) and symptomatic (4 months) stages across different muscle types (Soleus, Gastrocnemius), fiber types, and subcellular regions (subsarcolemmal, intermyofibrillar). Mitochondrial network structure was altered as early as 1 month of age, whereas density changes became widespread at 4 months in a fiber-type- and region-dependent manner. Alongside early lysosomal enlargement and late-stage spatial clustering, we observed a progressive, region-specific increase in mitochondria-lysosome contacts, most pronounced in the intermyofibrillar region. This quantitative framework provides a powerful tool for monitoring pathophysiology and evaluating therapeutic interventions in Pompe disease.

## Introduction

Pompe disease (glycogen storage disease type II; OMIM #232300) is an autosomal recessive lysosomal storage disorder caused by pathogenic variants in the GAA gene, which encodes acid α-glucosidase (GAA), the lysosomal hydrolase responsible for glycogen degradation [1–3]. Loss of GAA activity leads to abnormal accumulation of glycogen within lysosomes, predominantly in skeletal, respiratory, and cardiac muscle, ultimately resulting in progressive muscle weakness and atrophy [4,5]. Two primary clinical phenotypes are recognized. Infantile-onset Pompe disease (IOPD) is the most severe form, characterized by near-complete deficiency of GAA activity (<2% of normal), rapid disease progression, severe hypertrophic cardiomyopathy, and profound skeletal muscle hypotonia [6,7]. Late-onset Pompe disease (LOPD) is characterized by residual enzymatic activity (2-40%) and a broader clinical spectrum marked by slower but relentless progression, primarily affecting skeletal and respiratory muscles [4,7]. In both forms, skeletal muscle impairment is a defining clinical feature, driving progressive mobility loss and respiratory failure [8].

Histologically, affected myofibers display prominent vacuolation caused by the accumulation of enlarged, glycogen-filled lysosomes alongside extensive autophagic debris The extent of lysosomal pathology varies according to the metabolic profile of the individual myofiber, suggesting that fiber-type-specific metabolic properties modulate susceptibility to lysosomal impairment [9,10]. A hallmark of Pompe disease muscle pathophysiology is massive autophagic buildup, reflecting defective fusion between autophagosomes and dysfunctional lysosomes; this blockade of autophagic flux exacerbates myofiber structural damage and disease progression [9,11,12].

In healthy skeletal muscle, mitochondria form a highly organized, interconnected mosaic network divided into two functionally distinct subpopulations: subsarcolemmal (SS) mitochondria, located beneath the sarcolemma and primarily supporting membrane-associated energetic processes; and intermyofibrillar (IM) mitochondria, which are distributed between myofibrils to supply ATP directly to the contractile apparatus [13–15]. Maintaining this compartmentalized arrangement is essential for muscle bioenergetics and mechanical performance. Structural mitochondrial abnormalities, including fragmentation, paracrystalline inclusions, loss of cristae, tubular elongation, and marked swelling, have been widely documented in muscle biopsies from Pompe patients and in *Gaa*^⁻/⁻^ mice [16–18]. Furthermore, defects in mitochondrial membrane potential and impaired mitophagy have been demonstrated in cultured *Gaa*^⁻/⁻^ myoblasts [17]. Growing evidence indicates that organelle crosstalk is severely disrupted in Pompe disease, as autophagic aggregates create physical and functional barriers that impede inter-organelle communication [19].

To date, characterization of the mitochondrial compartment in Pompe disease has relied predominantly on transmission electron microscopy (EM) [16–19]. Although EM provides exceptional structural resolution, its specific limitations (small field of view, 3D reconstruction technically demanding, complex sample preparation, time and cost) hinder comprehensive evaluation of large-scale network connectivity and spatial organization across whole myofibers. Consequently, significant gaps remain regarding how mitochondria, depending on skeletal muscle fiber type (oxidative Type I vs. glycolytic Type II) and their subcellular region (SS vs. IM), undergo pathological remodeling over time, and how they physically interact with lysosomes and autophagosomes.

Super-resolution fluorescence microscopy techniques overcome the classical diffraction limit (∼200 nm) of confocal microscopy [20]. Stimulated emission depletion (STED) microscopy achieves lateral resolutions of ∼30 nm and can resolve mitochondrial cristae periodicity [21–23]. Single-molecule localization microscopy (SMLM) approaches such as STORM, DNA-PAINT, and PALM push lateral resolution down to ∼20 nm, albeit typically at the expense of field of view [20].

In the present study, we established an integrated methodology combining super-resolution fluorescence microscopy with deep learning–based image segmentation to quantitatively characterize mitochondrial mosaic architecture in skeletal muscle while simultaneously mapping its spatial interplay with lysosomal and autophagosomal compartments. We utilized image scanning microscopy (ISM), which achieves a lateral resolution of ∼130 nm while preserving a wide field of view, thereby optimizing the trade-off between spatial resolution and volumetric sampling coverage [24]. Unlike STED or SMLM, which typically require restricting the field of view to achieve their highest resolutions [20,25], mapping mitochondrial network architecture and organelle interactions across multiple intact myofibers requires broad spatial sampling, for which a moderate super-resolution enhancement over confocal microscopy is ideal. Applying this workflow to healthy wild-type (WT) and *Gaa*^-/-^ mice, we performed a quantitative multi-organelle analysis considering muscle metabolic composition (glycolytic Gastrocnemius vs. mixed oxidative/glycolytic Soleus), fiber type (Types I and II), subcellular mitochondrial location (SS vs. IM), and disease stage (pre-symptomatic 1-month vs. symptomatic 4-months). Our findings show that this imaging approach provides a powerful framework for visualizing and quantitatively analyzing interactions between organelles within skeletal muscle fibers, thus offering new opportunities to explore the subcellular changes underlying disease progression. As such, it represents an interesting tool for monitoring pathophysiology of a wide range of muscle diseases such as mitochondriopathies and evaluating efficacy of therapeutic strategies.

## Results

### Super-resolution imaging highlights muscle microenvironment-dependent mitochondrial alteration in Pompe disease mice

We first sought to determine the extent to which overall mitochondrial network distribution and architecture are altered in skeletal muscle during Pompe disease progression using super-resolution microscopy. Quantitative descriptors classically used to characterize mitochondrial network morphology (density, branching, and branch length [26,27]) were evaluated in longitudinal sections of Soleus and Gastrocnemius muscles from 1- and 4-month-old wild-type (WT) and *Gaa*^⁻/⁻^ mice. Because the Gastrocnemius muscle consists predominantly of fast-twitch glycolytic fibers, analysis was restricted to Type II fibers [28,29]. Conversely, the Soleus muscle, which exhibits a mixed oxidative/glycolytic profile, was analyzed by separating Type I and Type II myofibers [28,29].

Quantification was performed using a dedicated image analysis pipeline (Fig. 1 and Supplementary Fig. S1). Translocase of outer mitochondrial membrane 20 (TOMM20)-positive signals were segmented using a trained "Segment.ai" deep-learning model integrated into a General Analysis 3 (GA3) workflow. Segmented mitochondrial structures were assigned to individual myofibers. To distinguish between mitochondrial subpopulations, subsarcolemmal (SS) and intermyofibrillar (IM) regions were delineated by applying circular erosion to binary myofiber outlines and subtracting the eroded mask from original outlines. This procedure preserved the geometric fidelity of the fibers while enabling precise segregation and classification of SS and IM mitochondria for downstream analysis.

**Figure 1.**
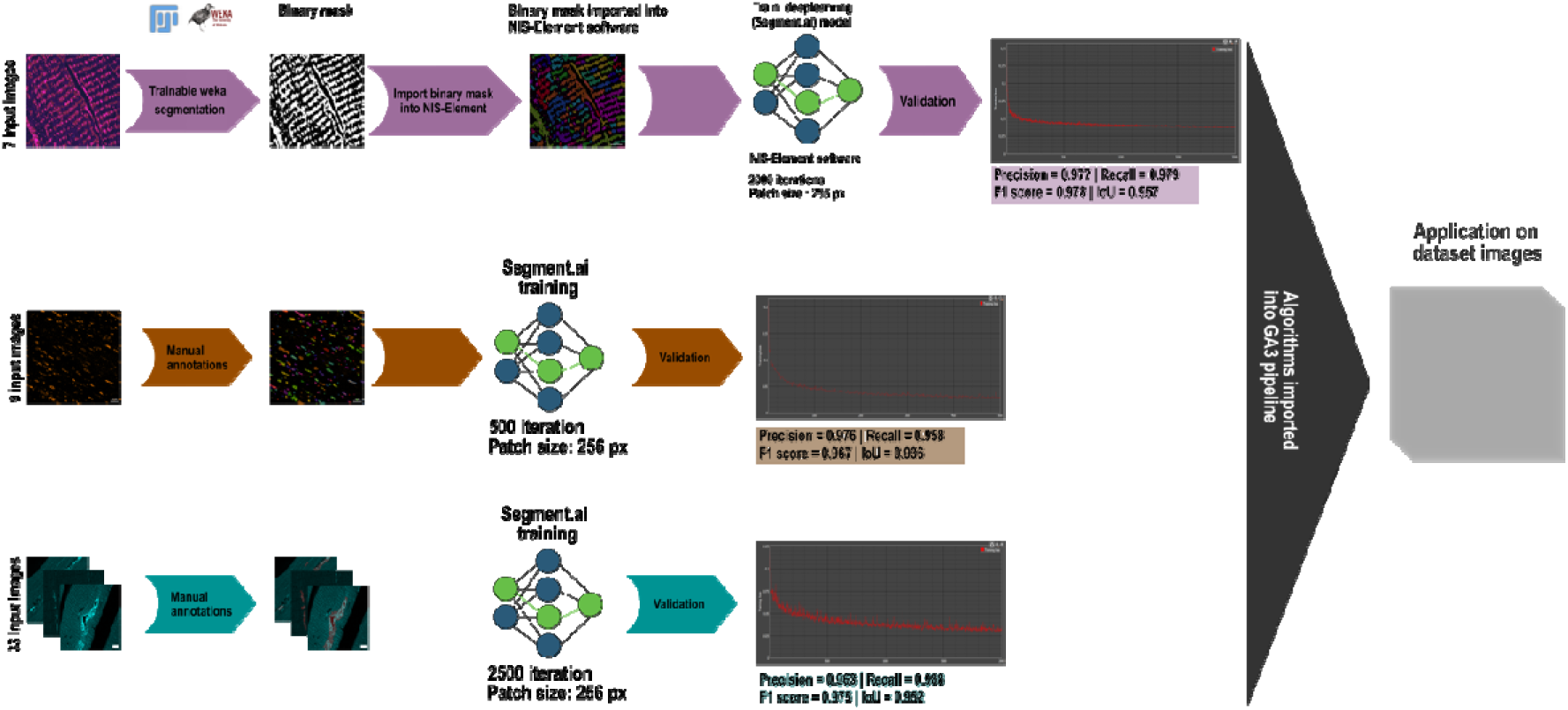
Deep-learning training strategies for organelle segmentation. Representative workflow for deep learning segmentation of mitochondria (TOMM20, magenta), lysosomes (LAMP-1, orange), and autophagosomes (LC3, sky blue). Cropped image datasets were manually annotated and used to train U-Net-based deep learning models ("Segment.ai") in NIS-Elements software, followed by model validation on an independent dataset of manually annotated ground-truth images

Super-resolution observation of TOMM20-labeled longitudinal sections revealed swollen mitochondrial structures in 1-month-old *Gaa*⁻/⁻ mice that were absent in WT controls at any age (Fig. 2A). Mitochondrial density, defined as the ratio of segmented mitochondrial surface area to total myofiber surface area, did not differ significantly between WT and *Gaa*⁻/⁻ mice at the pre-symptomatic stage (1 month), with the exception of the IM region in Soleus Type II fibers, revealing no major change in mitochondrial density at pre-symptomatic stage of the disease (p < 0.01; Fig. 2B; Table I). At 4 months of age, mitochondrial density in Soleus IM regions was significantly increased specifically in Type I fibers (p < 0.01). In the Gastrocnemius muscle, 4-month-old *Gaa*⁻/⁻ mice exhibited marked increases in mitochondrial density in both IM (p < 0.001) and SS (p < 0.01) regions of Type II fibers relative to WT controls.

**Figure 2.**
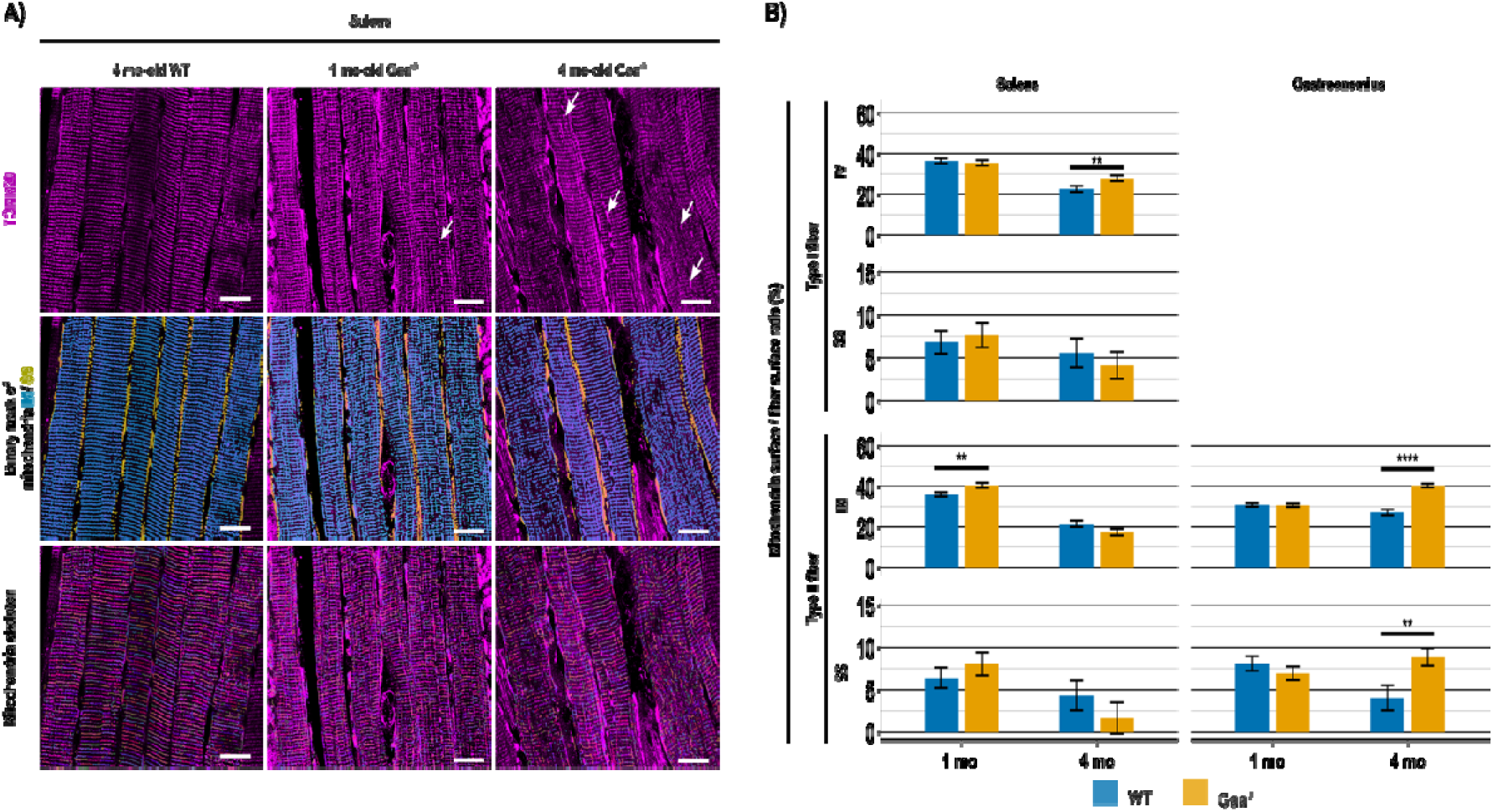
Super-resolution image scanning microscopy reveals muscle- and fiber-type-dependent mitochondrial remodeling in Pompe disease mice. (A) Representative super-resolution image scanning microscopy (ISM) images of mitochondrial networks in longitudinal Soleus muscle sections from wild-type (WT) and *Gaa*^⁻/⁻^ mice labeled with anti-TOMM20 (top row: raw fluorescence input; middle row: binary masks generated by deep learning segmentation; bottom row: skeletonized networks illustrating branching architecture). Arrowheads highlight swollen mitochondrial structures in *Gaa*^⁻/⁻^ muscle. Scale bar = 20 µm. (B) Quantitative analysis of mitochondrial surface density (mitochondrial surface area relative to total myofiber area, %) in Soleus (Type I and Type II fibers) and Gastrocnemius (Type II fibers) muscles of 1- and 4-month-old WT and *Gaa*^⁻/⁻^ mice, separated into intermyofibrillar (IM) and subsarcolemmal (SS) regions. Data are presented as mean ± SEM (*n* = 4 animals per group). Statistical analysis was performed using a linear mixed-effects model. ** p < 0.01, *** p < 0.001 vs. age-matched WT controls.

**Table I.**
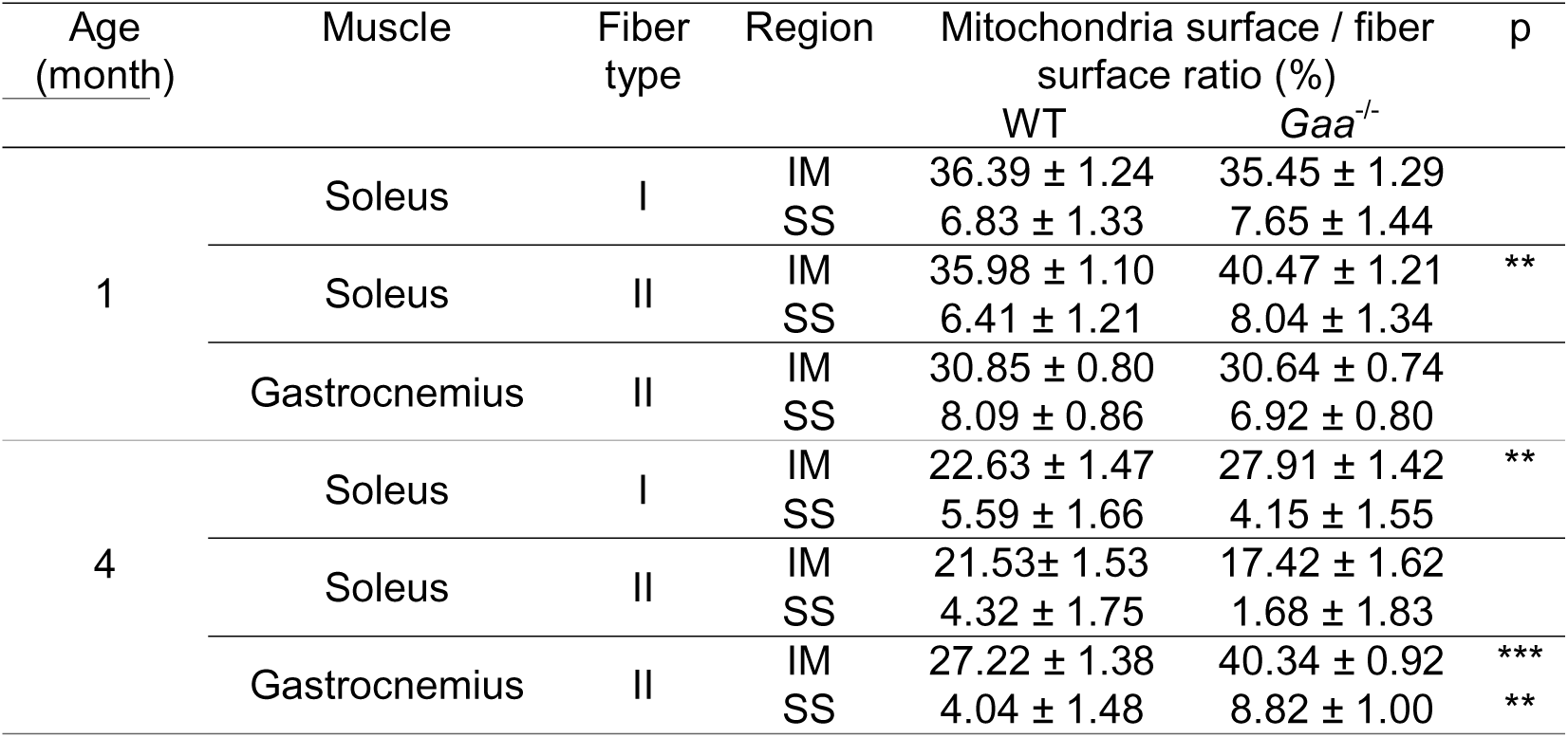
Mitochondrial surface density per myofiber area. Mitochondrial surface area expressed as a percentage of total myofiber surface area. Values represent mean ± SEM. SS: Subsarcolemmal; IM: Intermyofibrillar. ** p < 0.01, *** p < 0.001 vs. age-matched WT.

To further evaluate network architecture, binary mitochondrial masks were skeletonized to measure branch point density (number of branch points normalized to mitochondrial surface area) and average branch length. At 1 month of age, branch point density in *Gaa*⁻/⁻ mice was significantly lower than in WT controls in both Type I and Type II fibers of the Soleus muscle (p < 0.001), but was significantly higher in Type II fibers of the Gastrocnemius muscle (p < 0.01; Table II). By 4 months of age, branch point density in *Gaa*⁻/⁻ mice was markedly higher in Soleus Type I fibers (p < 0.001) and Gastrocnemius Type II fibers (p < 0.001), with a strong upward trend in Soleus Type II fibers (p = 0.054). Branch length differed between genotypes exclusively in Soleus Type II fibers, in which it was significantly higher at 1 month (p < 0.05) and significantly lower at 4 months (p < 0.05; Table II) of age.

**Table II.**
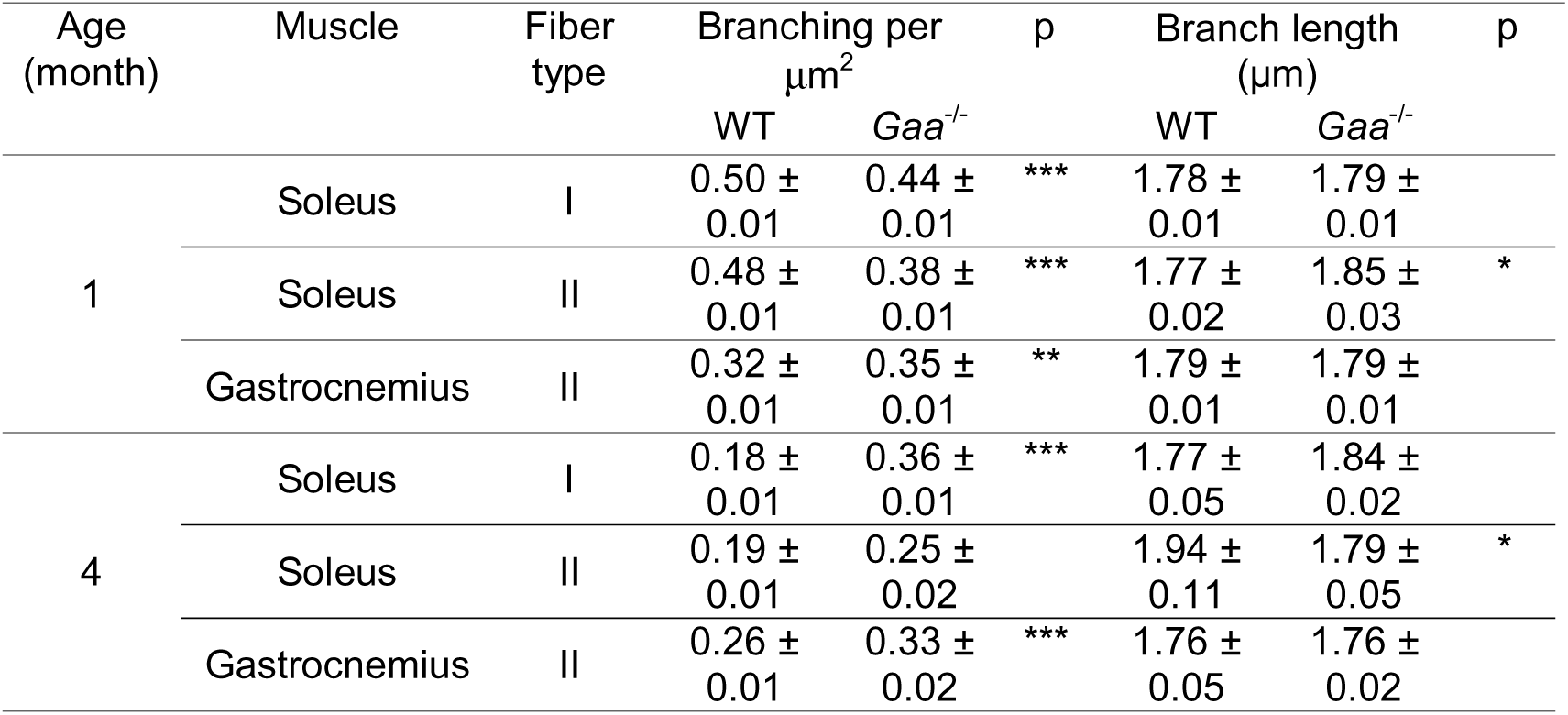
Mitochondrial branching point density and branch length. Branch point density expressed as number of branch points per mm² of mitochondrial surface area; branch length expressed in micrometers (µm). Values represent mean ± SEM. * p < 0.05, ** p < 0.01, *** p < 0.001 vs. age-matched WT.

Collectively, these data demonstrate that while mitochondrial density alterations become widespread at 4 months, structural remodeling of the mitochondrial network (branching and length) is evident as early as 1 month of age. These pathophysiological changes depend strongly on muscle type, fiber type, and subcellular location, revealing the presence of a complex, spatiotemporally regulated network reorganization during disease progression.

### Early lysosomal enlargement and spatial clustering in Pompe disease mouse muscle

To investigate whether mitochondrial network alterations correlate with primary lysosomes and autophagosome defects in Pompe disease, we next characterized lysosomal dimensions (LAMP-1⁺ objects) and autophagosomal surface occupancy (LC3⁺ objects) across SS and IM regions and myofiber types (Fig. 3A). Given previous reports showing that lysosomal storage varies according to myofiber metabolic profile [10,19], lysosomal Feret diameter and surface density (ratio of lysosomal surface area to total myofiber area) were quantified in Type I and Type II fibers.

**Figure 3.**
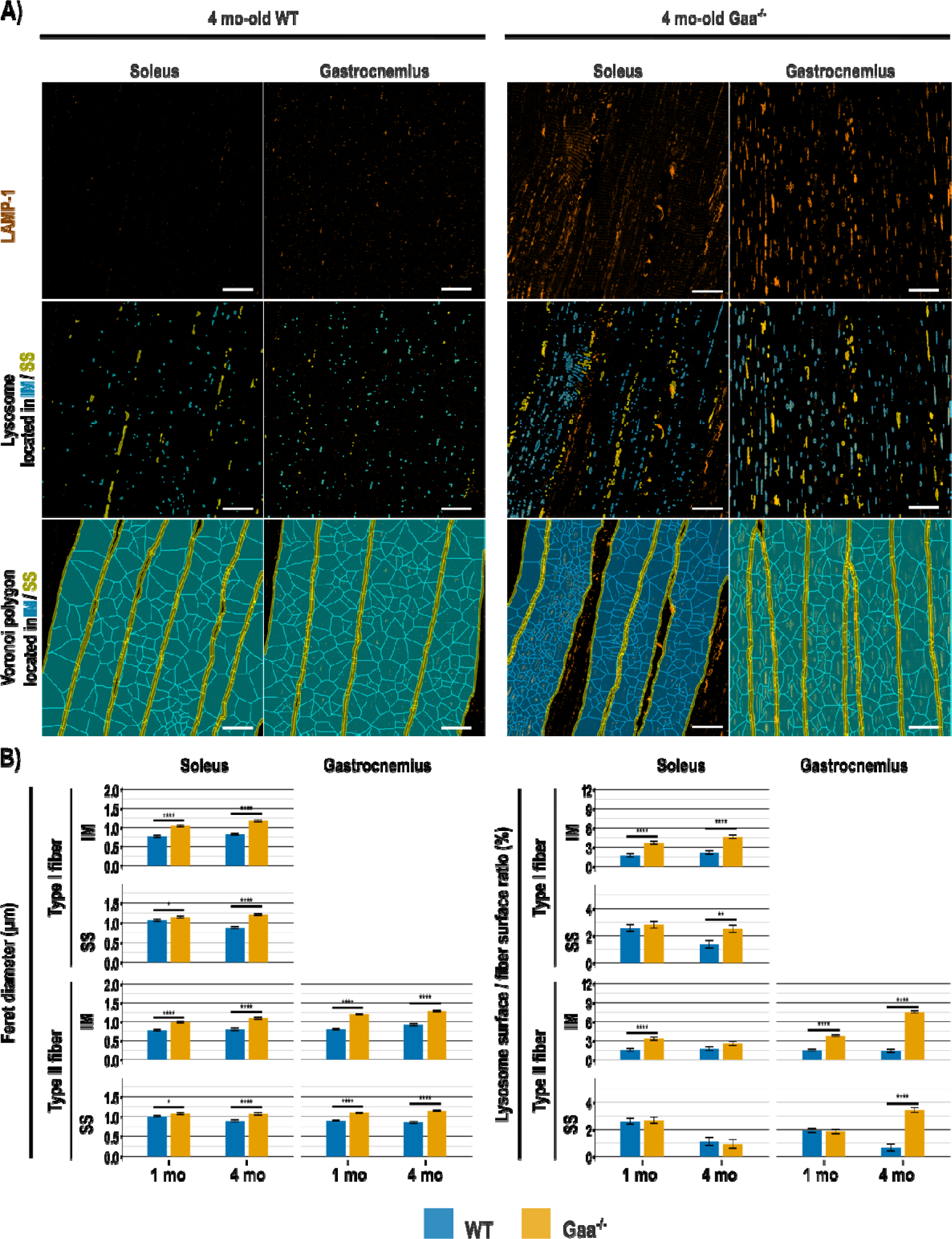
Early lysosomal enlargement and late-stage spatial lysosomal clustering in Pompe disease skeletal muscle. (A) Representative super-resolution ISM images of lysosomes in longitudinal muscle sections from wild-type (WT) and *Gaa*^⁻/⁻^ mice. Top row: raw fluorescence input (LAMP-1, orange); middle row: binary masks of segmented lysosomes; bottom row: Voronoi tessellation diagrams illustrating spatial distribution and proximity between neighboring lysosomes within myofibers. Scale bar = 20 µm. (B) Quantitative evaluation of lysosomal Feret diameter (µm) and surface density (lysosomal area relative to myofiber area, %) across subsarcolemmal (SS) and intermyofibrillar (IM) subregions in Soleus (Type I and Type II) and Gastrocnemius (Type II) fibers at 1 and 4 months of age. Data represent mean ± SEM (*n* = 4 animals per group). Statistical analysis was performed using a linear mixed-effects model. * p < 0.05, ** p < 0.01, *** p < 0.001, ****p < 0.0001) vs. age-matched WT controls.

Beginning at 1 month of age, lysosomal Feret diameter was significantly higher in *Gaa*⁻/⁻ mice compared with WT controls across all muscles, fiber types, and subcellular regions, confirming early, generalized lysosomal enlargement (p < 0.05 to p < 0.001; Fig. 3B; Table III). By contrast, higher lysosomal surface area at 1 month were restricted to IM regions of Type I and Type II Soleus fibers and Type II Gastrocnemius fibers (p < 0.001; Table III), indicating early region-specific lysosomal enlargement. By 4 months of age, lysosomal surface area in Soleus muscle was significantly increased in both IM (p < 0.001) and SS (p < 0.01) regions, but specifically in Type I fibers. In the Gastrocnemius muscle, lysosomal expansion was markedly amplified, with *Gaa*⁻/⁻ Type II fibers demonstrating a 5.3-fold higher IM surface density (p < 0.001) and a 5.2-fold higher SS surface density (p < 0.001) versus WT controls. These results point to distinct, fiber-type-dependent patterns of lysosomal alterations.

**Table III.**
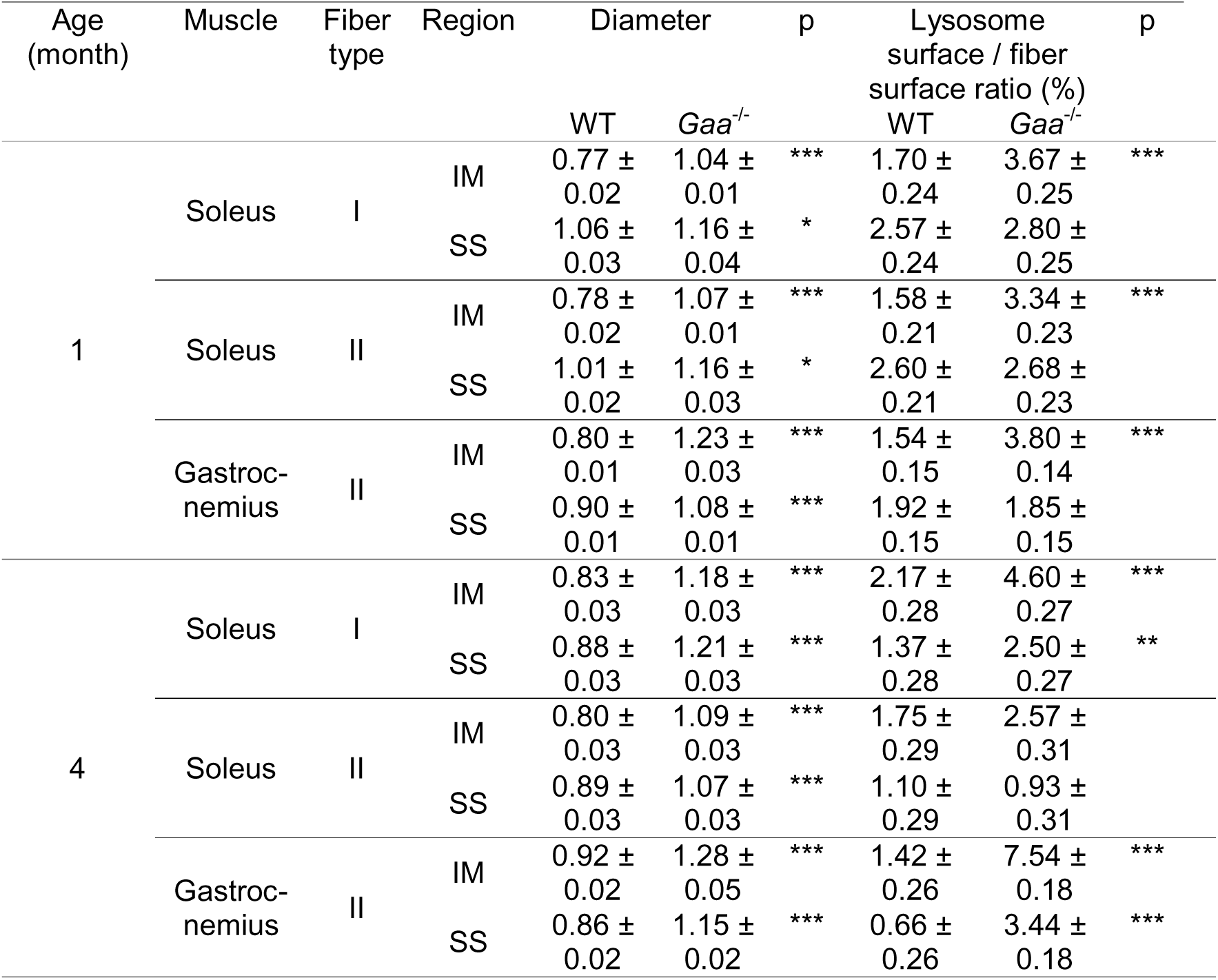
Lysosomal Feret diameter and surface density. Feret diameter expressed in micrometers (µm); lysosomal surface area expressed as percentage of myofiber area. Values represent mean ± SEM. IM: Intermyofibrillar; SS: Subsarcolemmal. * p < 0.05, ** p < 0.01, *** p < 0.001 vs. age-matched WT.

To evaluate whether lysosomal spatial distribution was altered, Voronoi polygon analysis was performed on binary lysosomal masks, using equivalent polygon diameter as an index of inter-lysosomal spacing (lower values indicate spatial clustering). At 1-month, inter-lysosomal distances were comparable between genotypes, except for a minor increase in the SS region of Gastrocnemius Type II fibers (p < 0.05; Table IV). By 4 months, inter-lysosomal distances were significantly lower in the IM region of Soleus Type I (p < 0.001), Soleus Type II (p < 0.001), and Gastrocnemius Type II (p < 0.001) fibers in *Gaa*⁻/⁻ versus WT mice, demonstrating progressive, tight lysosomal clustering. Within the SS region, these distances were significantly shorter only in type II fibers of the Gastrocnemius (p < 0.05) in *Gaa*⁻/⁻ versus WT mice.

**Table IV.** Lysosomal spatial distribution (Voronoi polygon equivalent diameter). Values expressed in micrometers (µm), representing inter-lysosomal distance (mean ± SEM). Lower values indicate tighter clustering. IM: Intermyofibrillar; SS: Subsarcolemmal. * p < 0.05, *** p < 0.001 vs. age-matched WT.

| Age (month) | Muscle | Fiber type | Region | Voronoi Equivalent Diameter ($\mu\text{m}$) | | p |
| --- | --- | --- | --- | --- | --- | --- |
|  |  |  |  | WT | *Gaa*<sup>-/-</sup> |  |
| 1 | Soleus | I | IM | 6.53 ± 0.16 | 6.18 ± 0.17 | |
| | | | SS | 2.75 ± 0.16 | 2.68 ± 0.16 | |
| | Soleus | II | IM | 6.36 ± 0.14 | 6.26 ± 0.15 | |
| | | | SS | 2.71 ± 0.14 | 2.57 ± 0.15 | |
| | Gastrocnemius | II | IM | 6.45 ± 0.10 | 6.66 ± 0.11 | |
| | | | SS | 2.68 ± 0.10 | 3.02 ± 0.11 | * |
| 4 | Soleus | I | IM | 7.24 ± 0.19 | 6.27 ± 0.18 | *** |
| | | | SS | 3.07 ± 0.19 | 2.83 ± 0.18 | |
| | Soleus | II | IM | 7.33 ± 0.20 | 6.15 ± 0.20 | *** |
| | | | SS | 3.05 ± 0.18 | 2.58 ± 0.18 | |
| | Gastrocnemius | II | IM | 7.65 ± 0.18 | 6.46 ± 0.12 | *** |
| | | | SS | 3.07 ± 0.17 | 2.62 ± 0.12 | * |

Finally, myofiber surface area occupied by autophagosomes (LC3⁺ objects) was significantly higher at 4 months than at 1 month of age in *Gaa*⁻/⁻ mice, an effect that was restricted to Type II fibers of the Gastrocnemius muscle (1.37 ± 0.08% vs. 0.90 ± 0.06%, p < 0.001; Supplementary Fig. S2). Together, these findings reveal early, widespread lysosomal enlargement in *Gaa*^-/-^ mice muscle fibers that is initially confined to the intermyofibrillar domain and subsequently spreads to the subsarcolemmal compartment in a muscle- and fiber-type-dependent manner.

### Early and markedly higher intermyofibrillar mitochondria-lysosome interactions in Pompe disease mice muscle

Given that mitochondria–lysosome contacts regulate key physiological processes including mitochondrial fission and calcium homeostasis [30,31], we next analyzed organelle physical crosstalk. Direct contact events, defined as border-to-border adjacency or embedment between segmented TOMM20⁺ and LAMP-1⁺ objects, were quantified across myofiber subregions (Fig. 4A).

**Figure 4.**
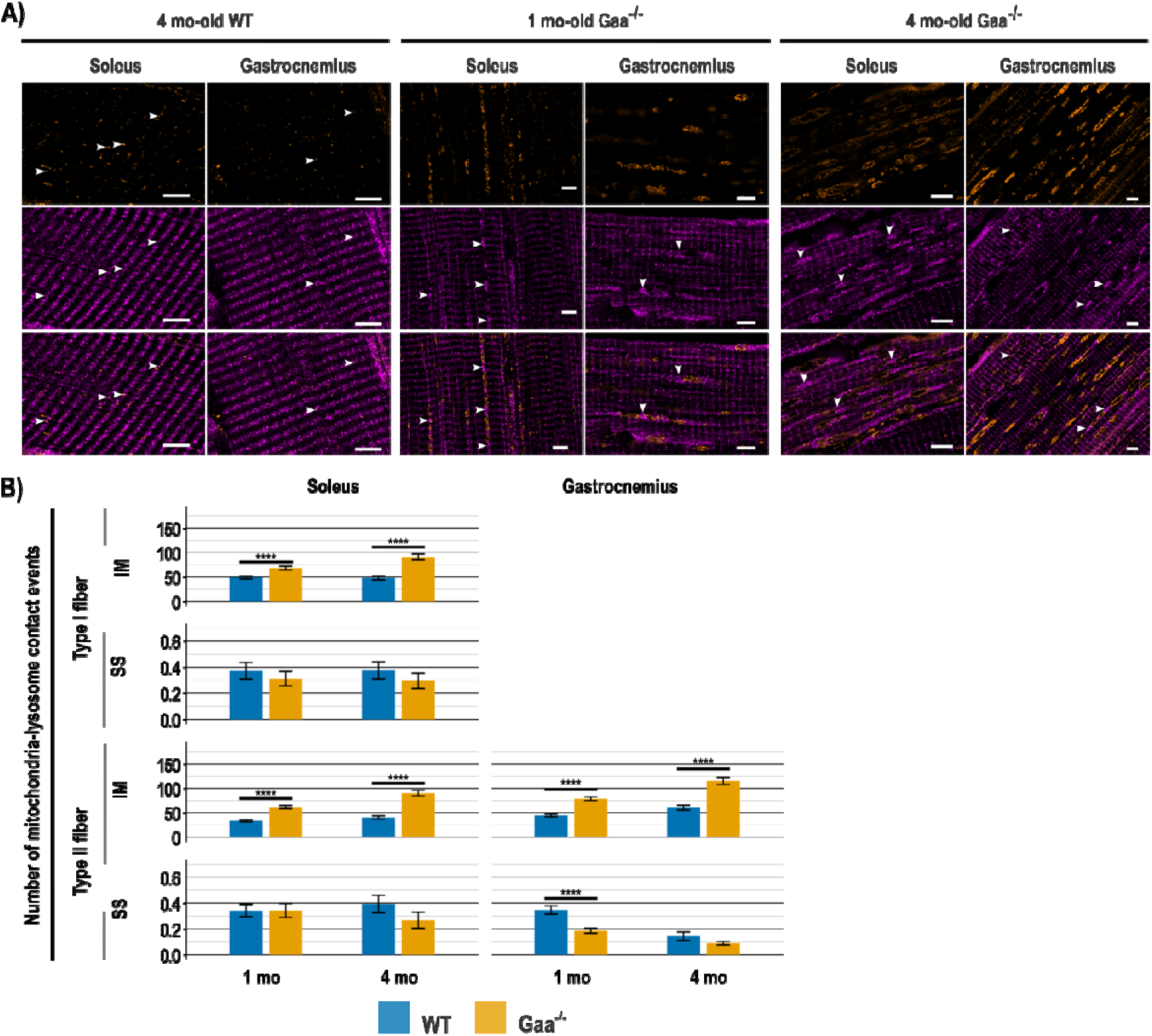
Elevated intermyofibrillar mitochondria–lysosome physical contact sites in Pompe disease muscle. (A) Representative super-resolution ISM images of mitochondria (TOMM20, magenta) and lysosomes (LAMP-1, orange) in longitudinal muscle sections from WT and *Gaa*^⁻/⁻^ mice. Top rows: single-channel images; bottom row: merged color overlay. Arrowheads highlight direct physical contact sites (<100 nm distance threshold) between mitochondria and lysosomes. In *Gaa*^⁻/⁻^ mice, altered mitochondria (arrowheads) were often closely associated with lysosomal structures. Scale bar = 5 µm. (B) Quantitative assessment of direct mitochondria–lysosome contact events per region in Soleus (Type I and Type II) and Gastrocnemius (Type II) fibers from 1- and 4-month-old WT and *Gaa*^⁻/⁻^ mice. Data represent mean ± SEM (*n* = 4 animals per group). Statistical analysis was performed using a linear mixed-effects model. *** p < 0.001, ****p < 0.0001 vs. age-matched WT controls.

At 1 month of age, the number of mitochondria–lysosome contact events in *Gaa*^⁻/⁻^ mice was significantly elevated in the IM region of Soleus Type I (*p* < 0.001) and Type II (*p* < 0.001) fibers, as well as Gastrocnemius Type II fibers (*p* < 0.001; Fig. 4B; Table V). By contrast, a small but significant reduction in contact events was observed in the SS region of Gastrocnemius Type II fibers (*p* < 0.001). By 4 months of age, the number of IM mitochondria–lysosome contact events was further increased across all examined muscle and fiber types (*p* < 0.001), with fold-increases in *Gaa*^⁻/⁻^ vs. WT ranging from 1.88- to 2.25-fold (compared to 1.38- to 1.84-fold at 1 month; Table V). Notably, in the SS region of Gastrocnemius Type II fibers, the number of contact events no longer differed significantly between genotypes at 4 months, indicating that the early SS reduction was transient.

**Table V.** Mitochondria–lysosome contact events. Quantification of the number of direct contact events per region (mean ± SEM). SS: Subsarcolemmal; IM: Intermyofibrillar. *** p < 0.001 vs. age-matched WT.

| Age<br>(month) | Muscle | Fiber<br>type | Region | Number of contact event<br>WT | *Gaa*<sup>-/-</sup> | p |
| --- | --- | --- | --- | --- | --- | --- |
| 1 | Soleus | I | IM | 49.17 ± 2.94 | 68.19 ± 3.70 | *** |
| | | | SS | 0.37 ± 0.06 | 0.31 ± 0.05 | |
| | Soleus | II | IM | 33.46 ± 1.99 | 61.43 ± 3.34 | *** |
| | | | SS | 0.34 ± 0.04 | 0.34 ± 0.05 | |
| | Gastrocnemius | II | IM | 44.32 ± 2.58 | 78.65 ± 4.13 | *** |
| | | | SS | 0.35 ± 0.03 | 0.19 ± 0.02 | *** |
| 4 | Soleus | I | IM | 48.34 ± 3.99 | 90.77 ± 5.94 | *** |
| | | | SS | 0.38 ± 0.06 | 0.30 ± 0.06 | |
| | Soleus | II | IM | 40.12 ± 3.32 | 90.19 ± 5.96 | *** |
| | | | SS | 0.39 ± 0.07 | 0.26 ± 0.06 | |
| | Gastrocnemius | II | IM | 60.27 ± 4.96 | 115.20 ± 7.45 | *** |
| | | | SS | 0.14 ± 0.03 | 0.09 ± 0.01 | |

## Discussion

Mitochondria are well-described as the "powerhouses" of the cell with essential roles in bioenergetics, calcium buffering, and diverse signaling pathways [32]. Their functional partnership with lysosomes is increasingly recognized as vital for the regulation of mitochondrial dynamics and calcium homeostasis [30,31]. In lysosomal storage disorders, mitochondrial dysfunction, characterized by organelle fragmentation, impaired respiration, and reduced ATP production, is a well-documented pathological feature [33]. In Pompe disease specifically, ultrastructural studies have reported swollen mitochondria, paracrystalline inclusions, loss of cristae, and fragmented morphology [16,19]. Subsequent *in vitro* studies in cultured myoblasts from *Gaa*^⁻/⁻^ mice confirmed mitochondria morphological changes and impaired metabolism [17]. However, there have been limited studies in skeletal muscle tissue that account for fiber-type metabolic heterogeneity, disease progression over time, and spatial organelle crosstalk.

In this study, we combined super-resolution image scanning microscopy (ISM; lateral resolution ∼130 nm) with deep learning-based segmentation to systematically map mitochondrial and lysosomal pathology in Pompe disease skeletal muscle *in situ*. By integrating organelle morphometrics with fiber-type identity, subcellular location, and longitudinal timepoints, our methodology enables precise detection of mitochondria, lysosomes, and autophagosomes within skeletal muscle, moving beyond qualitative description to establish a quantitative framework for analyzing disease progression.

Consistent with previous ultrastructural studies in patients and animal models [16,18,19], our findings show that structural remodeling of the mitochondrial network begins prior to clinical symptom onset and precedes widespread changes in organelle density. Swollen mitochondria and decreased branching density were detected in *Gaa*^⁻/⁻^ muscle at 1 month of age. By contrast, changes in mitochondrial surface density were initially confined to a single subpopulation (intermyofibrillar [IM] region of Soleus Type II fibers) at 1 month, becoming more widespread across muscles and regions at the later, symptomatic stage. This progressive pattern suggests that mitochondrial density changes are, at least in part, secondary to lysosomal expansion and autophagic build-up, which could physically disrupt mitochondrial positioning and connectivity as storage accumulates. We hypothesize that early branching alterations reflect compensatory structural dynamics, whereas late-stage density shifts are secondary to progressive lysosomal expansion and autophagosome accumulation, which physically displace and disorganize the mitochondrial network. Furthermore, skeletonization analysis revealed branch length differences between *Gaa*^-/-^ and WT mice specifically in Soleus Type II fibers, with a transient increase at 1 month followed by a significant decrease at 4 months. These data, which reflect a reorganization of the mitochondrial compartment, are consistent with those reported in cultured myoblasts from 5- to 6-month-old *Gaa*^⁻/⁻^ mice [17] and provide additional information on the complexity of mitochondrial network remodeling.

In the lysosomal compartment, alterations were evident from the earliest post-natal stage examined (1 month), consistent with its primary role in disease pathogenesis. Lysosomes showed a significant increase in diameter across all muscle subregions at 1 month of age, and formed compact clusters at 4 months of age, an effect that was particularly pronounced in the Gastrocnemius muscle [9,10,19]. This progressive transition from isolated enlarged lysosomes to dense spatial clusters induces severe cytoplasmic crowding, imposing mechanical and metabolic stress on neighboring organelles and the contractile apparatus. These dense spatial clusters of lysosomes observed at later disease stages imposes mechanical and metabolic stress on the surrounding cytoplasm, thereby exacerbating myofiber dysfunction. Based on these findings, we demonstrate that the analytical pipeline developed here provides a comprehensive quantitative framework to characterize lysosomal remodeling, overcoming the limitations of purely descriptive analyses.

A key finding of the present study is the identification of pronounced alterations in mitochondria-lysosome interactions as a pathological hallmark of Pompe disease in skeletal muscle. Specifically, the number of contact sites was higher, elevated from 1 month of age within the IM region of Soleus (Types I and II) and Gastrocnemius (Type II) fibers, with even greater differences evident by 4 months. This preferential involvement of IM mitochondria in contact formation may reflect their close proximity to contractile elements and increased energetic demands. Because mitochondria-lysosome contacts have been implicated in mitochondrial quality control in physiological conditions [30,31], we hypothesize that their greater abundance and altered spatial distribution in Pompe disease may become maladaptive, potentially contributing to mitochondrial dysfunction, impaired mitophagy, and energetic imbalance [17,19,34].

Taken together, these results demonstrate the feasibility of applying deep learning-based segmentation to resolve complex subcellular structures, such as mitochondria, in skeletal muscle tissue. Moreover, these findings are consistent with previous studies in which early lysosomal enlargement and clustering precede, and may contribute to, subsequent alterations in mitochondrial network architecture, suggesting a possible link between lysosomal dysfunction and mitochondrial impairment [12,17,19].

Enzyme replacement therapy (ERT), the current standard of care for Pompe disease, yields incomplete correction of skeletal muscle pathology [8,35,36]. As such, the quantitative, multi-organelle framework presented here could serve as a valuable platform to assess whether novel therapies normalize mitochondrial network architecture and mitochondria-lysosome crosstalk alongside glycogen clearance. Extending this study to more advanced stages of the disease would allow to refine the proposed sequence of organelle remodeling. Our analysis was performed on longitudinal sections, which allow visualization of organelle organization along the fiber axis but are limited to a thin layer at an uncontrolled depth within the fiber volume. Also, such analysis restrict quantification to fibers parallel to the section plane. Preparations of isolated fibers would allow us to overcome these two limitations and gain access to the complete 3D organization of mitochondria and lysosomes. Finally, validation in human biopsy material will be an important next step to assess the translational relevance and generalization of these findings.

In conclusion, our integrated super-resolution imaging and analytical pipeline establishes a robust framework for quantifying mitochondrial network architecture and organelle crosstalk *in situ*. This methodology offers broad utility for investigating diseases involving mitochondrial dysfunction and provides a sensitive tool for evaluating therapeutic efficacy in Pompe disease.

## Materials and Methods

### Animals

All animal experiments were conducted in accordance with the ARRIVE guidelines (https://arriveguidelines.org) and European Council directives on the care and use of laboratory animals. Experimental protocols were reviewed and approved by the Ethics Committee on Animal Experimentation of the Pays de la Loire region, France (authorization APAFIS #43113).

*Gaa*^⁻/⁻^ mice (GAA-KO 6^neo^/6^neo^ model) harboring a targeted disruption of exon 6 in the *Gaa* gene, resulting in complete loss of acid α-glucosidase enzymatic activity, were utilized. Heterozygous breeding pairs were originally provided by Dr. Nina Raben (National Institutes of Health, Bethesda, MD, USA) and maintained in a specific pathogen-free (SPF) animal facility at Oniris (Nantes, France). Homozygous *Gaa*^⁻/⁻^ mice and wild-type (WT) littermate controls were generated by homozygous and heterozygous breeding schemes, respectively. Animals were housed under controlled environmental conditions (22 °C; 12:12 h light:dark cycle) with *ad libitum* access to standard rodent chow and water. Environmental enrichment (nesting rolls) was provided to minimize distress and chronic stress. For each age group (1 and 4 months) and genotype (WT and *Gaa*^⁻/⁻^), *n* = 4 male animals were analyzed, from which Soleus and Gastrocnemius muscles were harvested.

### Tissue Sample Preparation

Mice were deeply anesthetized via intraperitoneal injection of a mixture of ketamine (75 mg/kg; Imalgene, Merial, Lyon, France) and xylazine (10 mg/kg; Rompun, Bayer, Leverkusen, Germany). Depth of anesthesia was confirmed by the absence of pedal withdrawal reflexes prior to euthanasia by intracardial perfusion. The thoracic cavity was opened, and phosphate-buffered saline (PBS) was perfused through the apex of the left ventricle, followed by systemic perfusion with 4% paraformaldehyde (PFA) in PBS. Soleus and Gastrocnemius muscles were then dissected, post-fixed overnight in 4% PFA at 4 °C, dehydrated through a graded ethanol series, cleared in xylene, and embedded in paraffin. Timepoints of 1 month (pre-symptomatic stage) and 4 months (onset of clinical symptoms) were evaluated.

### Histological and immunofluorescence analysis

Paraffin-embedded muscle samples were sectioned at an 8 µm thickness using a microtome and mounted on glass slides. Slides were dried overnight at 37 °C, deparaffinized in xylene (3 × 10 min; MC3007581000, VWR, Radnor, PA, USA), and rehydrated through a graded ethanol series in distilled water: 100% ethanol (3 × 10 min), 95% ethanol (10 min), 70% ethanol (10 min), and 50% ethanol (10 min). Sections were then transferred to PBS. Heat-induced antigen retrieval was performed in Heat-induced antigen retrieval was performed using preheated Tris-EDTA buffer (pH 9.0; ZUC029, Zytomed, Berlin, Germany) in a water bath at 98 °C for 40 min, followed by cooling at room temperature (RT) for 20 min.

Sections were washed in PBS and permeabilized with 0.2% Triton X-100 (X100, Sigma-Aldrich, Saint-Quentin-Fallavier, France) in PBS for 30 min at RT. Non-specific binding was blocked by incubation for 30 min at RT in blocking buffer containing 10% goat serum, 2% bovine serum albumin (BSA; A7906, Sigma-Aldrich), and 0.2% Triton X-100 in PBS. Serial cross-sections were incubated overnight at 4 °C with primary antibodies diluted in blocking buffer. Primary antibodies included: anti-TOMM20 (rabbit monoclonal, 1:200; ab186734, Abcam, Cambridge, UK); anti-citrate synthase (rabbit monoclonal, 1:200; MA517264, Thermo Fisher Scientific, Waltham, MA, USA) to label mitochondria; anti-LAMP-1 (rat monoclonal, 1:50; 553792, BD Biosciences, Franklin Lakes, NJ, USA) to label lysosomes; anti-LC3 (rabbit monoclonal, 1:100; 83506, Cell Signaling Technology, Danvers, MA, USA) to label autophagosomes. Following PBS washes (3 × 5 min), sections were incubated for 1 h at RT with species-appropriate secondary antibodies: goat anti-rabbit IgG Alexa Fluor 488 (1:300; A21131, Invitrogen, Carlsbad, CA, USA); goat anti-rat IgG Alexa Fluor 555 (1:300; A21434, Invitrogen); goat anti-rabbit IgG Alexa Fluor 647 (1:400; A21245, Invitrogen); polyclonal goat anti-rabbit DyLight 405 (1:300; 115-475-207, Jackson ImmunoResearch, West Grove, PA, USA).

Sections were washed in PBS and mounted using ProLong Gold antifade mounting medium (11559306, Thermo Fisher Scientific).

For myofiber typing, adjacent longitudinal sections were incubated for 1 h at 37 °C with primary antibodies against slow myosin heavy chain (sMHC/Type I; rabbit monoclonal, 1:400; ab234431, Abcam) and fast myosin heavy chain (fMHC/Type II; mouse monoclonal, 1:500; M4276, Sigma-Aldrich). After PBS washes, sections were incubated for 1 h at RT with goat anti-rabbit Alexa Fluor 555 (1:800; A21428, Invitrogen) and goat anti-mouse Alexa Fluor 647 (1:1000; A21240, Invitrogen). Nuclei were counterstained with DAPI (1:2000 in distilled water, 10 min; 10184322, Thermo Fisher Scientific), washed, and mounted in ProLong Gold antifade mounting medium.

### Fiber-typing imaging

Whole-section myofiber type imaging was conducted using an AxioScan slide scanner (Zeiss, Jena, Germany) equipped with an XCite LED FIRE illumination system (Excelitas Technologies, Waltham, MA, USA), a Plan-Apochromat 10× objective (NA = 0.45), and an ORCA-Flash4.0 monochrome sCMOS camera (Hamamatsu Photonics, Hamamatsu, Japan).

### Super-resolution imaging of organelles

Super-resolution fluorescence imaging was performed on a laser scanning confocal AX-NSPARC microscope system (Nikon Europe BV, Amstelveen, Netherlands). Sequential channel excitation was employed to prevent spectral crosstalk: the first pass excited Alexa Fluor 488, followed by subsequent passes for remaining fluorophores. Emission fluorescence was collected using a 502-546 nm bandpass filter and a quad-band filter set (430-663 nm / 582-618 nm / 663-691 nm). A Plan-Apochromat 60× oil immersion objective (NA = 1.42) was used.

Field of view (FOV) dimensions were set to 150 × 150 µm and sampled at 2048 × 2048 pixels, yielding an isotropic pixel size of 70 nm. Each FOV was positioned to encompass at least five longitudinal myofibers. For quantitative sampling, a minimum of 5 regions of interest (ROIs) for Soleus and 10 ROIs for Gastrocnemius were acquired per muscle. Image datasets were deconvoluted using the Richardson-Lucy algorithm with 15 iterations prior to feature extraction.

### Deep learning model training and segmentation

Segmentation of TOMM20 (mitochondria), LAMP-1 (lysosomes), and LC3 (autophagosomes) signals was executed using a U-Net–based deep learning model, "Segment.ai," implemented in NIS-Elements software (v6.10; Nikon Europe BV).

#### TOMM20 model

Seven cropped XY image stacks were manually annotated using the Trainable Weka Segmentation plugin in Fiji/ImageJ. Annotations were imported into NIS-Elements, and the model was trained for 2,000 iterations on an NVIDIA RTX5000 graphics processing unit (GPU) (driver v516.40; NVIDIA, Santa Clara, CA, USA).

#### LAMP-1 model

Nine cropped image stacks were manually annotated directly in NIS-Elements and trained for 500 iterations on the same GPU architecture.

#### LC3 model

Thirty-three cropped image stacks representing diverse morphological variants and signal-to-noise ratios were annotated and trained for 2,500 iterations.

Models were validated against an independent ground-truth dataset of 10 manually annotated images. Segmentation performance was evaluated using Precision (proportion of correct predictions), Recall (true positive rate), F1 score (harmonic mean of Precision and Recall), and Intersection over Union (IoU / Jaccard Index) (measures overlap between the predicted segmentation mask and the ground truth mask).

### Image analysis pipeline

Quantitative feature extraction was executed using the General Analysis 3 (GA3) module in NIS-Elements (v6.10.1; Nikon Europe BV). The GA3 workflow (Supplementary Fig. S1) was applied to 417 ROIs encompassing 2,640 individual myofibers.

Fiber segmentation and subregion delineation: Binary masks of muscle fibers were imported. To segregate subsarcolemmal (SS) from intermyofibrillar (IM) organelle populations, circular morphological erosion was applied to fiber masks. Subtracting eroded masks from original outlines defined the SS boundary while preserving geometric fiber outlines; remaining internal areas were assigned as IM regions.

Mitochondrial morphometrics: Segmented TOMM20 masks were assigned to SS or IM regions. Mitochondrial density was calculated as total segmented mitochondrial surface area divided by corresponding myofiber area. Mitochondrial binary masks were skeletonized to compute branch point density (number of branch points normalized to mitochondrial surface area) and individual branch length distributions. Lysosomal and autophagosomal morphometrics: Segmented LAMP-1 masks provided lysosomal surface density (lysosomal area / fiber area) and Feret’s maximum diameter. Spatial distribution was analyzed by generating Voronoi tessellation diagrams on binary lysosomal centroids, taking equivalent polygon diameter as a metric of inter-lysosomal proximity. Autophagosomal abundance was calculated as LC3-positive surface area relative to total fiber area. Organelle interactions: direct physical contact sites between mitochondria and lysosomes were scored when the distance between segmented TOMM20 and LAMP-1 boundary surfaces was <100 nm (border-to-border contact or embedment).

### Statistical analysis

Statistical computing was performed in R (v4.4.1). Continuous variables are expressed as mean ± standard error of the mean (SEM). To account for the hierarchical structure of nested data (multiple myofibers nested within individual animals), linear mixed-effects models were implemented using the lme4 package. Genotype (WT vs. *Gaa*^⁻/⁻^) was modeled as a fixed effect, with random intercepts assigned for individual animals to control for intra-animal correlation. Model assumptions, including independence and homoscedasticity/normality of residuals, were systematically verified using diagnostic plots. Statistical significance was established at p < 0.05.

## DECLARATIONS

### Ethics Approval and Consent to Participate

All experimental procedures involving animals were conducted in accordance with European Council guidelines and were reviewed and approved by the Ethics Committee on Animal Experimentation of the Pays de la Loire region, France (permit authorization APAFIS #43113).

### Consent for Publication

Not applicable.

### Availability of Data and Materials

The datasets generated and/or analyzed during the current study are available from the corresponding author upon reasonable request.

### Author Contributions

I.H. conceived, designed, and performed the experiments, analyzed the data, and wrote the manuscript. J.D. participated in experimental design, collected tissue samples, and sectioned muscle tissues. C.T. supervised statistical analysis, reviewed, and edited the manuscript. T.F. supervised the study and assisted with image acquisition. L.D. supervised the study, contributed to experimental design, data interpretation, and manuscript drafting. K.R. participated in study design, supervised the work, contributed to data interpretation, and edited the manuscript. M.A.C. conceived and supervised the study, contributed to data interpretation, and revised the manuscript. All authors read and approved the final manuscript.

## Acknowledgements

We thank the technical staff of the Oniris rodent facility (Nantes, France) for animal care. We are also grateful to the Nikon Imaging Centre at Institut Curie - CNRS for providing access to advanced microscopy equipment and expertise. We thank the APEX platform (INRAE/Oniris, Center of Excellence Nikon Nantes [CENN], Nantes, France) from UMR 0703 PAnTher (INRAE/Oniris, Nantes, France) for their valuable technological support. The authors also thank Biogenouest (the network of technology core facilities in life sciences and the environment in Western France, supported by the Conseil Régional des Pays de la Loire) for supporting UMR 0703 PAnTher and its APEX platform. We also gratefully acknowledge financial support from NeurATRIS: A Translational Research Infrastructure for Biotherapies in Neurosciences, “Investissement d’Avenir-ANR-11-INBS-0011” and from the Association Française contre les Myopathies (AFM) #23775.

## List of Abbreviations

ARRIVE: Animal Research: Reporting of *In Vivo* Experiments
BSA: Bovine serum albumin
DNA-PAINT: DNA points accumulation for imaging in nanoscale topography
EM: Transmission electron microscopy
ERT: Enzyme replacement therapy
fMHC: Fast myosin heavy chain
FOV: Field of view
GA3: General Analysis 3
GAA: Acid α-glucosidase (protein)
IM: Intermyofibrillar
IoU: Intersection over Union (Jaccard Index)
IOPD: Infantile-onset Pompe disease
ISM: Image scanning microscopy
LAMP-1: Lysosome-associated membrane protein 1
LC3: Microtubule-associated protein 1A/1B-light chain 3
LOPD: Late-onset Pompe disease
PALM: Photoactivated localization microscopy
PBS: Phosphate-buffered saline
PFA: Paraformaldehyde
ROI: Region of interest
sMHC: Slow myosin heavy chain
SMLM: Single-molecule localization microscopy
SS: Subsarcolemmal
STED: Stimulated emission depletion
STORM: Stochastic optical reconstruction microscopy
TOMM20: Translocase of outer mitochondrial membrane 20
WT: Wild-type

## Supplemental Figures

**Supplementary Figure 1 (Fig.S1).**
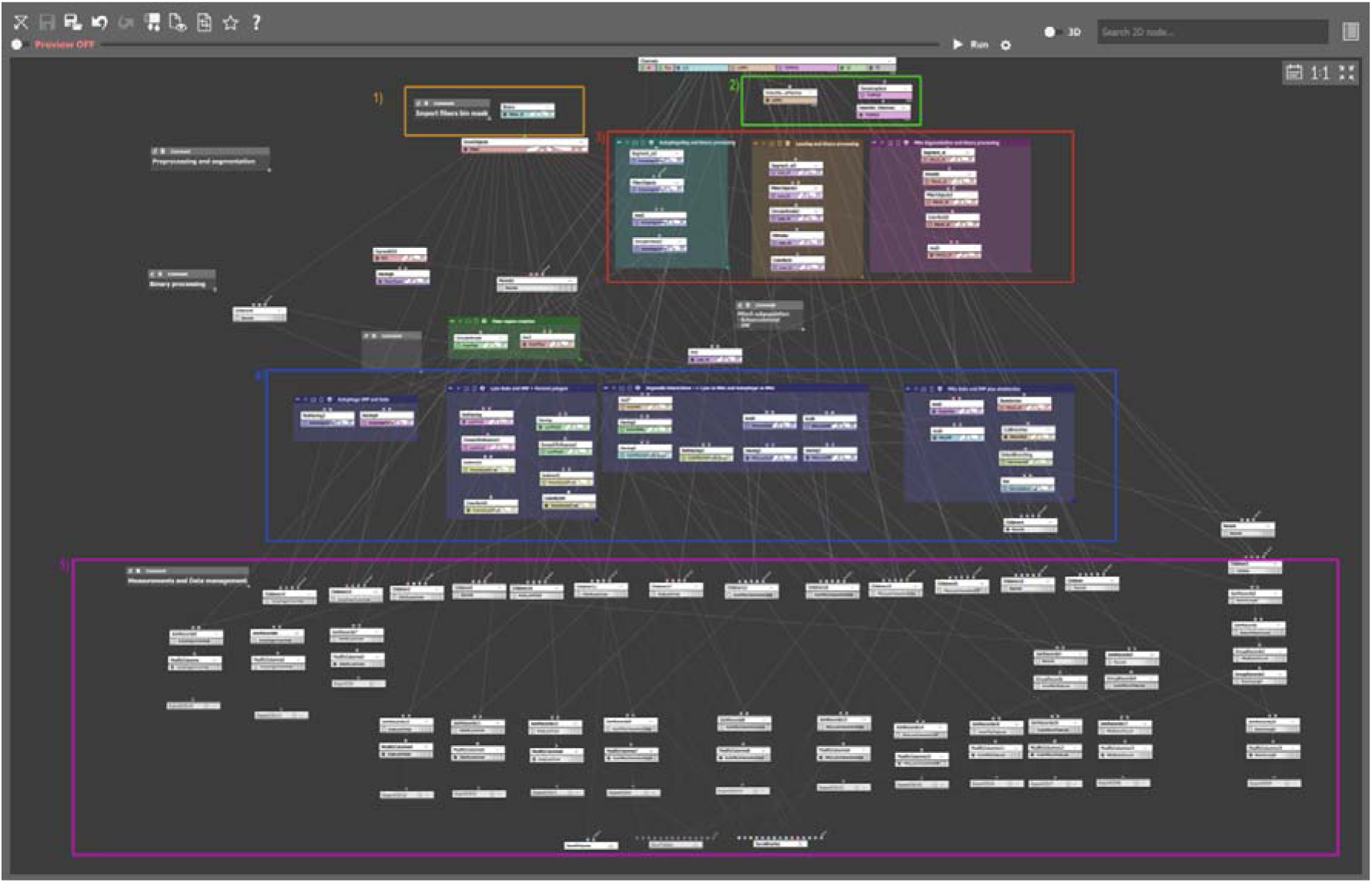
General Analysis 3 (GA3) pipeline for automated multi-organelle feature extraction. Detailed schematic of the GA3 analysis workflow developed in NIS-Elements for automated quantification of organelle morphometrics and physical interactions:1. Import and preprocessing (orange module): Binary masks of annotated muscle fibers were imported into the execution environment. 2. Contrast enhancement (green module): Local maxima contrast enhancement algorithms were applied to LAMP-1 and TOMM20 fluorescent channels to optimize organelle boundary detection. 3. Segmentation (red module): Three trained U-Net deep learning models ("Segment.ai") executed segmentation of autophagosomes, lysosomes, and mitochondria, followed by binary morphological smoothing. 4. Binary operations & spatial assignment (blue module): Myofibers were partitioned into subsarcolemmal (SS) and intermyofibrillar (IM) subregions using circular mask erosion. Organelles were assigned to their respective cellular compartments. Voronoi diagrams were constructed on lysosomal centroids to measure inter-lysosomal spatial spacing. Mitochondrial masks were skeletonized to calculate branch point density and branch length. Direct contact sites (<100 nm boundary proximity) between mitochondria and lysosomes/autophagosomes were mapped. 5. Quantifications (magenta module): Structural metrics (surface density, Feret diameter, branching density, branch length) and inter-organelle contact event counts were extracted per myofiber and subregion.

**Supplementary Figure 2 (Fig.S2).**
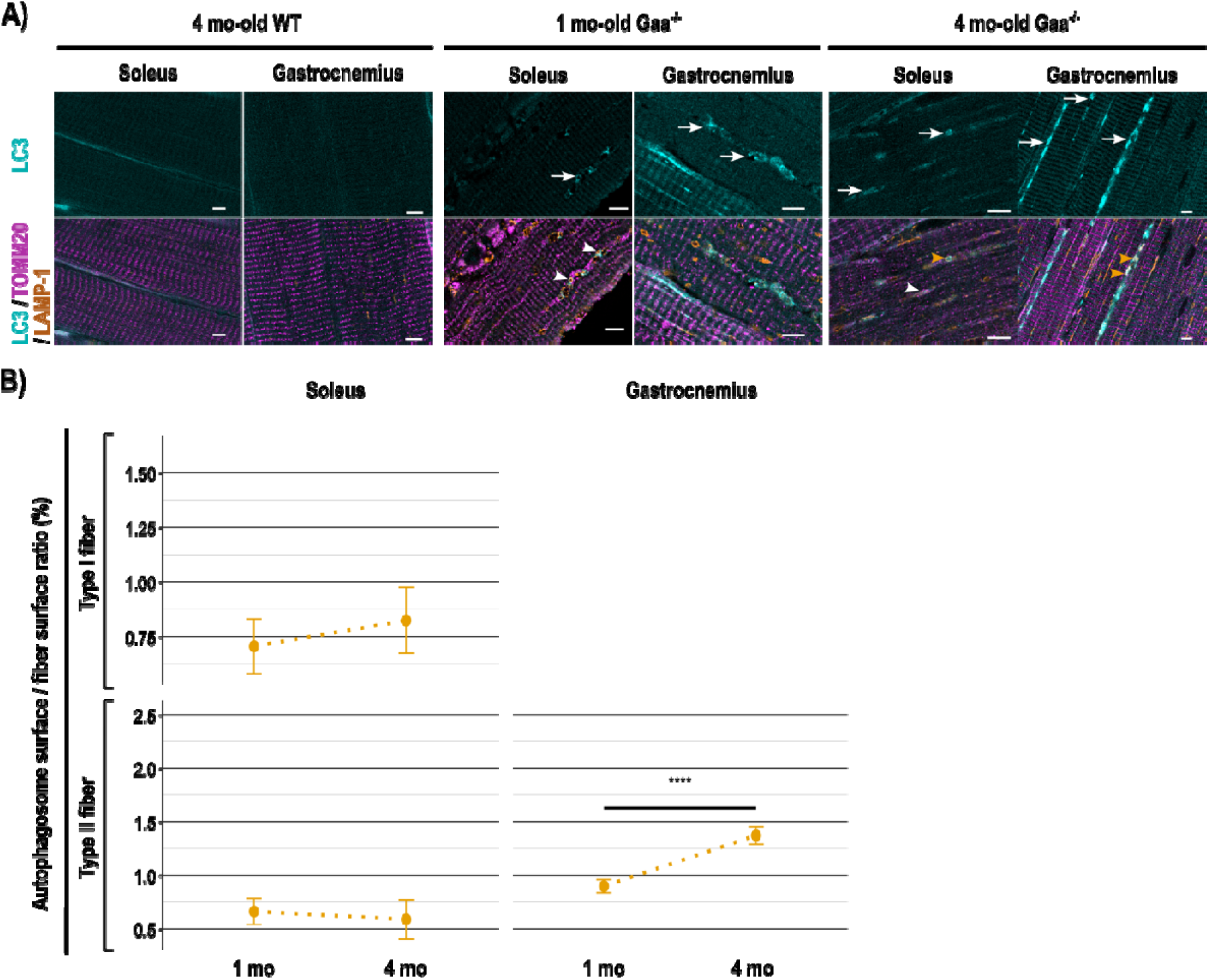
Autophagosome distribution and fiber-type surface occupancy in Pompe disease muscle. (A) Representative super-resolution ISM images of autophagosomes in longitudinal muscle sections. Top row: single-channel LC3 (cyan); bottom row: triple overlay showing LC3 (cyan), LAMP-1 (lysosomes, orange), and TOMM20 (mitochondria, magenta). In *Gaa*^⁻/⁻^ muscle, autophagic buildup is evident (white arrows). Autophagosomes are frequently observed adjacent to lysosomes alone (orange arrowheads) or simultaneously contacting both lysosomes and mitochondria (white arrowheads). Scale bar = 5 µm. (B) Quantitative surface occupancy of autophagosomes (ratio of LC3-positive surface area to total myofiber area, %) in Soleus (Type I and Type II) and Gastrocnemius (Type II) fibers from 1- and 4-month-old *Gaa*^⁻/⁻^ mice. Data represent mean ± SEM (*n* = 4 animals per group). Statistical analysis was conducted using a linear mixed-effects model. *** p < 0.001.

